# DNA-targeting and cell-penetrating antibody-drug conjugate

**DOI:** 10.1101/2023.04.12.536500

**Authors:** Anupama Shirali, Valentina Dubljevic, Fei Cao, Robert N. Nishimura, Allen Ebens, James A. Campbell, James E. Hansen

## Abstract

DNA released by dying cancer cells offers a tumor targeting strategy that is independent of specific cell surface antigens. Anti-DNA antibodies preferentially localize to tumor microenvironments enriched in extracellular DNA and can penetrate live tumor cells through nucleoside salvage pathways. Nuclear-localizing variants of anti-DNA antibodies cause DNA damage and selectively kill cancer cells with defects in DNA repair. Here we show that an optimized full-length IgG1 anti-DNA antibody penetrates live cells and is synthetically lethal to BRCA2-deficient tumors but has minimal effect of DNA repair-proficient tumors. Linkage of the antibody to the anti-mitotic drug monomethyl auristatin E yields a DNA-targeting and cell-penetrating anti-DNA antibody-drug conjugate (ADC) that is well tolerated in mice and highly toxic to tumors with intact DNA repair. This work provides proof-of-concept for the novel use of an anti-DNA antibody as the backbone of a DNA-targeting, cell-penetrating ADC that can impact tumors that otherwise lack specifically targetable surface antigens.

**Statement of significance:** A strategy for targeting tumors that lack specific surface antigens is revealed by an anti-DNA antibody-drug conjugate that localizes to tumor microenvironments enriched in DNA and penetrates cells through nucleoside salvage pathways.

## Introduction

DNA released by dying cancer cells offers the opportunity to leverage the biology of anti-DNA antibodies against malignancy. Anti-DNA antibodies associated with systemic lupus erythematosus exhibit preferential localization to tumor microenvironments enriched in extracellular DNA, and some are taken into live tumor cells through the activity of nucleoside salvage pathways. Inside tumor cells, nuclear-localizing variants of anti-DNA antibodies cause DNA damage and selectively kill cancer cells with pre-existing defects in DNA repair. Discovery of these anti-DNA antibody functions led to the design of the Deoxymab class of biologics composed of nuclear-localizing and DNA-damaging anti-DNA antibodies that are in preparation for clinical trials. (1-10)

The Deoxymabs are based in part on the prototype murine anti-DNA antibody 3E10 isolated from the MRL/lpr lupus mouse model. 3E10 exhibits DNA-dependent tumor localization, cellular penetration mediated by the nucleoside transporter ENT2, and selective killing of cancer cells with defects in DNA repair by causing lethal DNA double-strand breaks (DSBs) (4-7, 9, 10) to accumulate. An optimized fragment of 3E10 (Deoxymab-1, DX1) with optimized complementarity-determining regions (CDRs), reduced molecular weight, and lacking an Fc crosses the blood-brain barrier to target and suppress orthotopic brain tumors, including glioma and metastases, and is being advanced towards clinical trials (9).

In addition to single agent effects on DNA repair-deficient cancer cells, the Deoxymabs offer a potential platform for delivering cargoes to tumors. Antibody-drug conjugates (ADCs) targeting tumor surface antigens are an important part of therapeutic regimens in malignancies such as HER2+ breast cancer (11, 12), but are not applicable to tumors that lack such specific surface antigens. In contrast, tumor targeting by Deoxymabs does not require such surface antigen, which raises the possibility for their use in the design of a tumor agnostic Deoxymab ADC that seeks out DNA in the tumor microenvironment.

Previous success with 3E10 as a molecular delivery agent supports the feasibility of Deoxymab ADCs. Surface conjugation of 3E10 fragments to drug-loaded nanocarriers promoted accumulation of nanocarriers at tumors independent of surface antigen. Moreover, this strategy established an autocatalytic positive feedback cycle wherein increased tumor cell death caused by delivery of cargo drug caused the release of more DNA into the microenvironment, and consequently recruited additional 3E10-linked nanocarriers to the area (8). While this approach holds significant clinical potential for transport of nanocarriers to the tumor environment for drug release, the large size of the nanocarriers may interfere with mechanisms of cellular penetration by 3E10 or the Deoxymabs. Conversely, we hypothesized the small size of directly linked drugs in an ADC would be permissive of cellular penetration by a Deoxymab ADC.

DX1 is well suited to single agent therapy with its reduced size and absence of the Fc, but these features are limiting in the design of a Deoxymab ADC due to their effects on circulating half-life and sequence available for conjugation to drug. In the present study we designed, generated, and characterized Deoxymab-3 (DX3), a full-length IgG1 anti-DNA antibody based on the DX1 CDR sequences. DX3 characterization studies confirmed it retains the core Deoxymab traits of DNA binding, nuclear penetration, and selective toxicity to tumors with defects in DNA repair. Further, cell-penetrating activity was preserved when DX3 was linked to the anti-mitotic drug monomethyl auristatin E (MMAE). When tested in mice bearing breast cancer xenografts, the DX3 ADC yielded significantly greater tumor suppression compared to an isotype control ADC, establishing proof-of-concept for Deoxymab ADCs.

## Results

### DX3 binds DNA and penetrates cells

DX3 was designed as an IgG1 with DX1 CDR sequences and a Asn297Asp (N297D) mutation to reduce FcγR interactions. DX3 was produced in CHO cells and purified as previously described (9), and retention of DNA binding confirmed using a 30-mer single-stranded DNA oligonucleotide ligand in surface plasmon resonance (SPR) studies with the Biacore T200. Duplicate studies determined consistent KDs of 113 and 112 nM (**Fig. 1A**). DNA binding by the prototype 3E10 antibody directly correlates with its efficiency of cellular penetration (7). Consistent with this, cell penetration and nuclear localization by DX3 was confirmed in DLD1 colon cancer cells (**Fig. 1B**) and MCF7 breast cancer and U87 glioma cells (**Fig. 2**).

**Figure 1.**
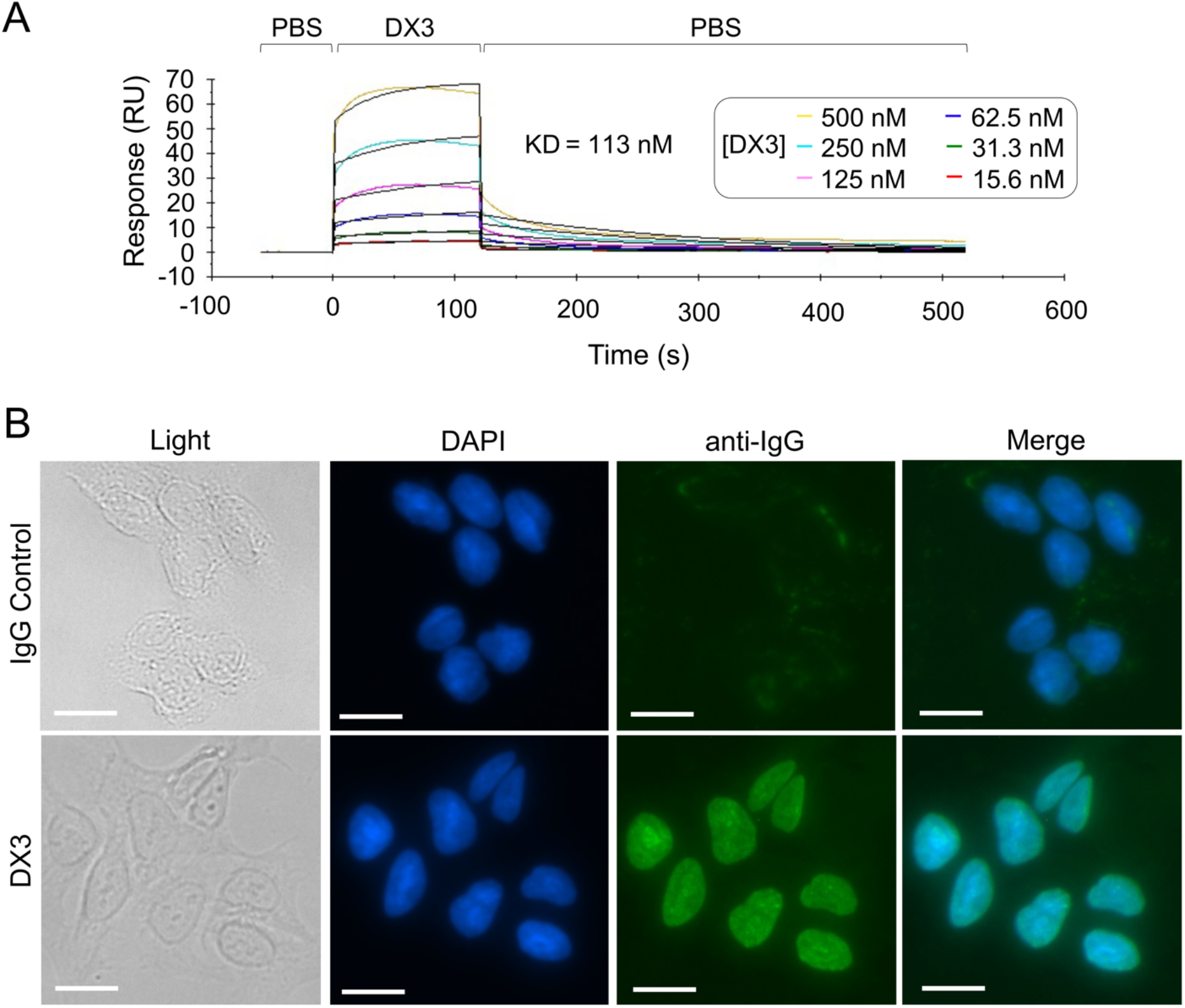
DX3 binds DNA and penetrates live cell nuclei. **(A)** DX3 binds single-stranded DNA with KD 113 nM. SPR binding profiles to 30-mer DNA oligonucleotide with DX3 titrated from 500 nM are shown. (**B**) DX3 penetrates live cell nuclei. Images of DLD1 colon cancer cells treated with 4 μM IgG control or DX3 and stained to detect antibody penetration by AF488 anti-IgG immunofluorescence with DAPI nuclear counterstain are shown. Bars: 20 μm.

**Figure 2.**
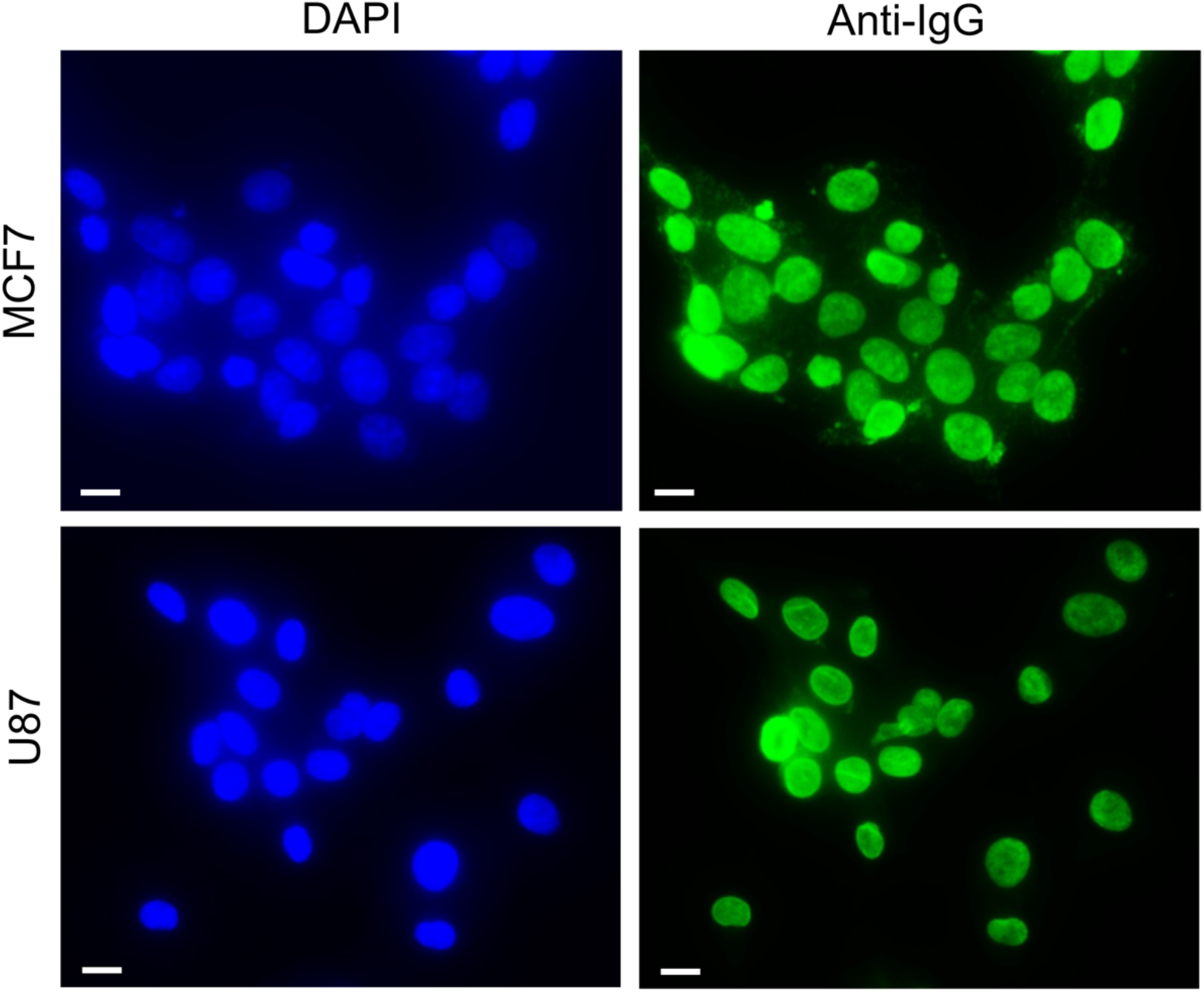
DX3 penetrates MCF7 and U87 cancer cells. Representative images of MCF7 breast cancer and U87 glioma cells treated with 10 μM DX3 and stained to detect antibody penetration by AF488 anti-IgG immunofluorescence with DAPI nuclear counterstain are shown. Bars: 20 μm.

### DX3 selectively suppresses BRCA2-deficient tumor growth

DNA binding and nuclear localization are defining features of the Deoxymabs, along with a third core trait of selective toxicity to tumor cells with defects in mechanisms of DNA repair, such as homologous recombination (HR). The impact of DX3 on the growth of HR-proficient and deficient tumors was examined using a matched pair of BRCA2-proficient and deficient DLD1 colon cancer cells, which were previously used in the development of DX1 (5, 6). Bilateral DLD1 flank tumors were generated by subcutaneous injection in nude mice, with BRCA2 WT tumors in the left flank and BRCA2-deficient in the right flank. When tumors reached mean volumes of ∼100 mm^3^, mice were randomized to treatment with vehicle control (N=8) or DX3 (50 mg/kg) (N=8) by tail vein injection twice a week. DX3 had no significant effect on the growth of BRCA2 WT tumors, with median tripling times of 9 days in mice treated with vehicle control or DX3, respectively (P=ns) (**Fig. 3A, B**). In contrast, DX3 significantly suppressed the growth of the contralateral BRCA2-deficient flank tumors, with tripling times of 9 days and not reached in mice treated with vehicle control or DX3, respectively (P=0.02) (**Fig. 3C, D**).

**Figure 3.**
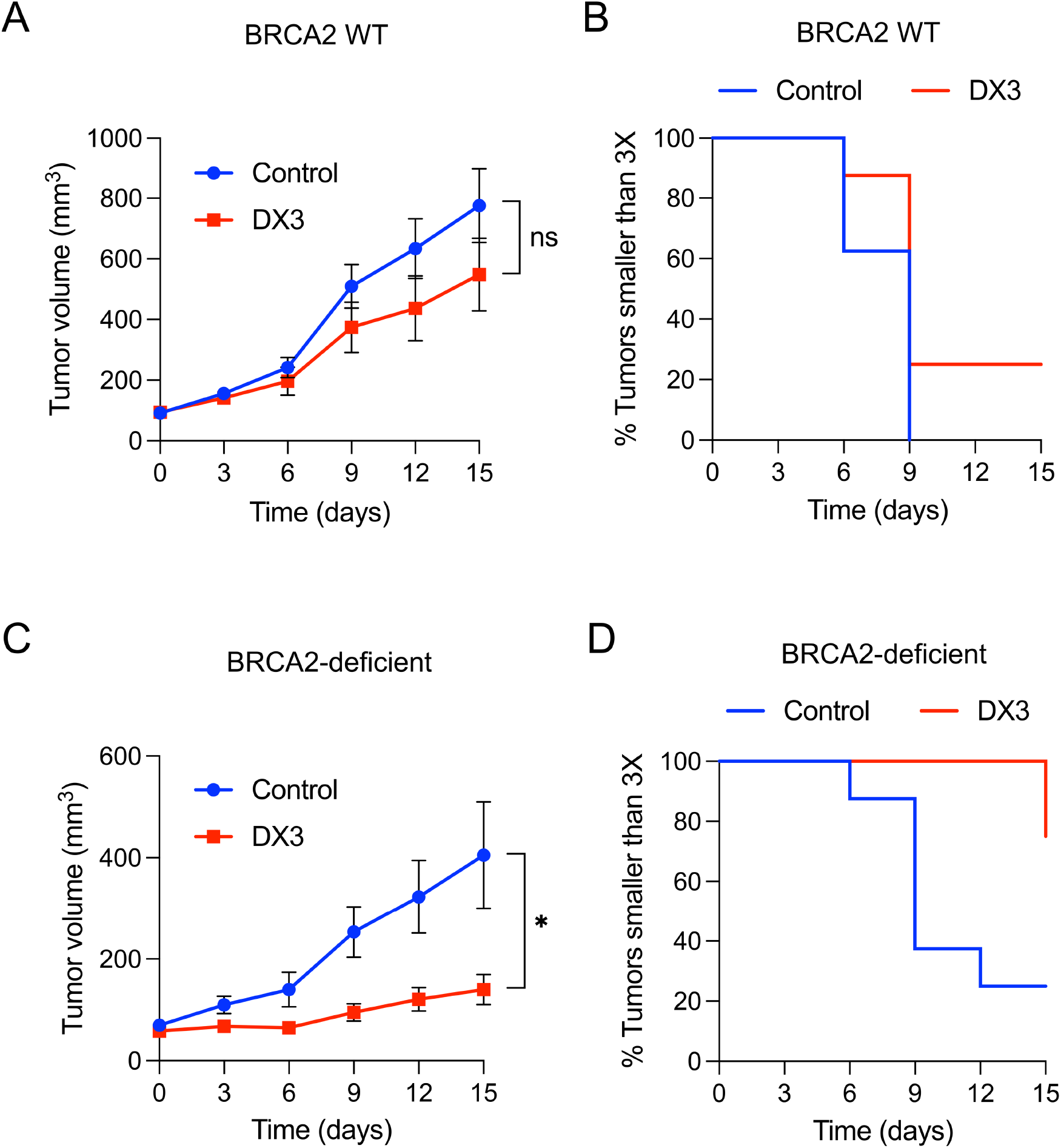
DX3 suppresses BRCA2-deficient tumor growth. (**A-D**) Tumor volumes and Kaplan-Meier plots of tumor tripling times in nude mice bearing bilateral DLD1 flank tumors (left flank WT, right flank BRCA2-deficient) treated with control or DX3 are shown. DX3 did not significantly impact BRCA2 WT tumor growth (P=ns, two-tailed student’s t-test) (**A**), with median tripling times of 9 days in mice treated with control or DX3 (P=ns, log-rank test) (**B**). In contrast, DX3 significantly suppressed the contralateral BRCA2-deficient tumors (*P<0.05, two-tailed student’s t-test) (**C**), with median tripling times of 9 days and not reached in mice treated with control or DX3, respectively (P=0.02, log-rank test) (**D**).

### Generation of a Deoxymab ADC

The Deoxymabs have single agent activity against DNA repair-deficient tumors but minimal effect on malignancies with intact mechanisms of repair. We hypothesized that a Deoxymab ADC that localizes and delivers cargo drug to tumors would be able to impact tumors with functional DNA repair. The anti-mitotic drug MMAE has been frequently used in ADC formulations, such as the CD30-targeting ADC brentuximab vedotin composed of an IgG1 linked to MMAE with a cleavable valine-citrulline (VC) linker (13). This drug and linker strategy were applied to DX3 to form DX3 vedotin. Briefly, DX3 was reduced with Tris (2-carboxyethyl phosphine (TCEP) followed by conjugation at room temperature to vcMMAE. Trastuzumab and an isotype control IgG1 were similarly conjugated to vcMMAE for use as control ADCs. To confirm purity and absence of free drug the resulting ADCs were evaluated by size exclusion high performance liquid chromatography (SEC-HPLC) and liquid chromatography mass spectrometry (LC-MS) and were >98% pure with undetectable free drug. Drug-antibody-ratio (DAR) in DX3 vedotin was 4.8 by hydrophobic interaction chromatography (HIC) and 4.5 by LC-MS (**Fig. 4**), closely matching the control ADC DARs of 4.7-4.9.

**Figure 4.**
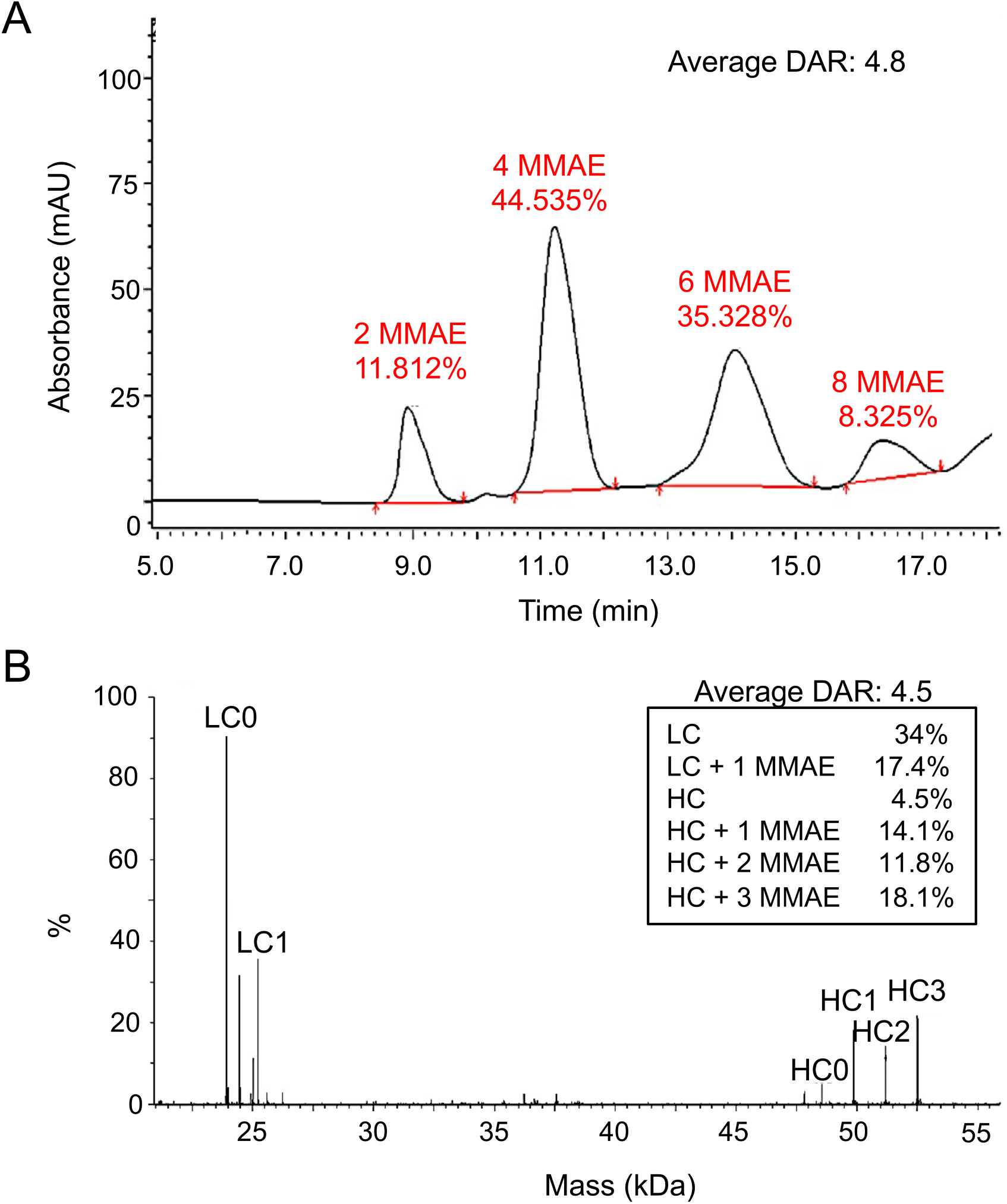
Determination of DX3 vedotin DAR. **(A)** DX3 vedotin HIC profile shows mean DAR 4.8, with four linked MMAE molecules the predominant species. (**B**) DX3 vedotin LC-MS profile shows mean DAR 4.5, consistent with the HIC results.

### DX3 vedotin retains cell-penetrating and MMAE functionality

U87 glioma and MCF7 breast cancer cells treated with trastuzumab vedotin or DX3 vedotin were immunostained to detect antibody penetration. While no significant cellular penetration by trastuzumab vedotin was observed, DX3 vedotin efficiently penetrated cells and localized into nuclei (**Fig. 5**). This result demonstrates that MMAE conjugation with a VC linker does not interfere with the efficiency of DX3 cellular penetration.

**Figure 5.**
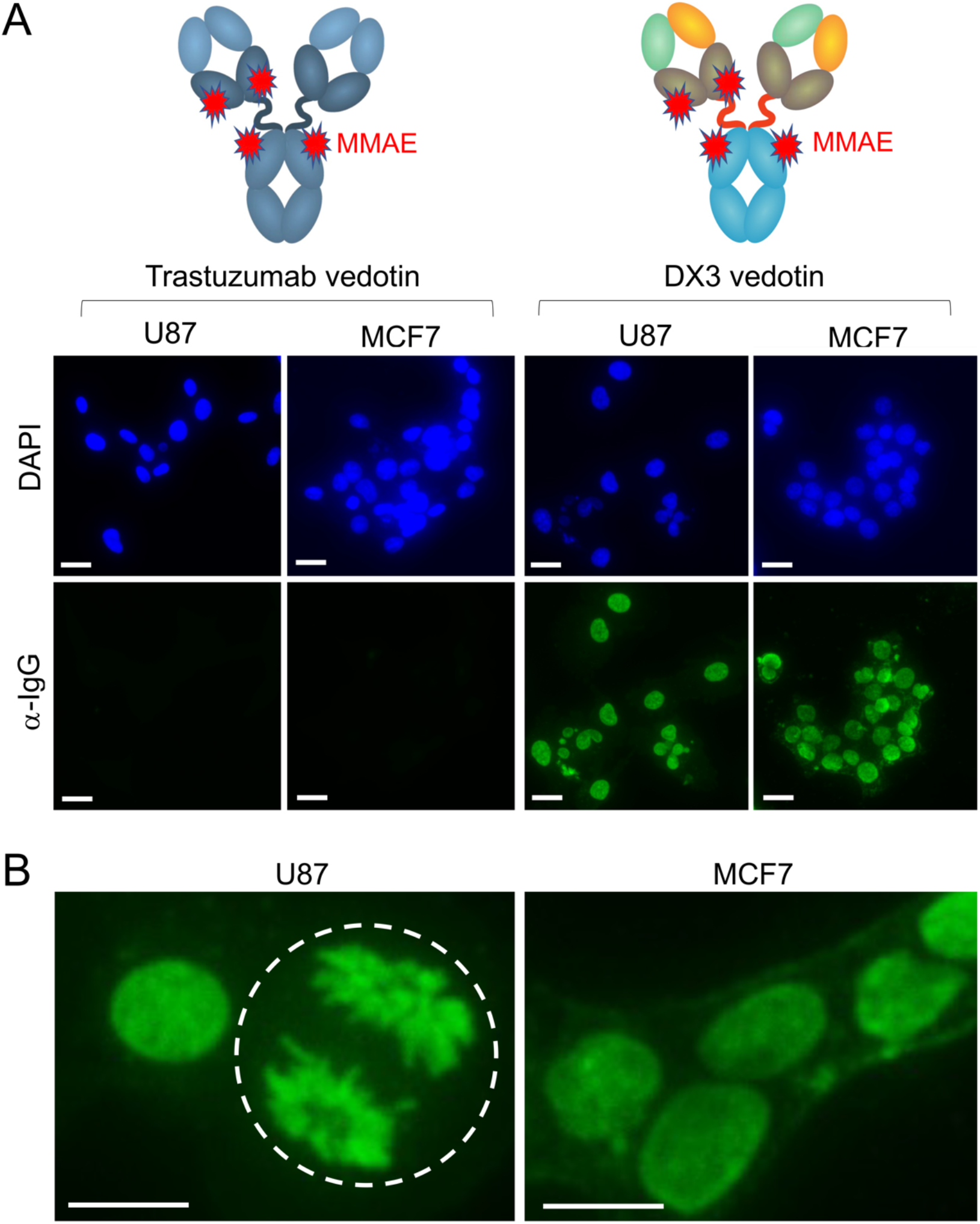
DX3 vedotin penetrates live cell nuclei. **(A and B)** Representative images of cells treated with 3 μM trastuzumab vedotin or DX3 vedotin and subsequently stained for antibody penetration by AF488 anti-IgG immunofluorescence with DAPI nuclear counterstain are shown in (**A**), with higher magnification images of DX3 vedotin-treated cells in (**B**). DX3 vedotin localized into the nuclei of both cell lines. DX3 vedotin association with chromosomes was readily seen in a cell in anaphase, denoted by the dashed circle in (**B**). Bars: 30 μm (**A**) and 15 μm (**B**).

Cells treated with MMAE exhibit rounding and membrane blebbing. Morphologies of U87 and MCF7 cancer cells treated with control media, DX3, trastuzumab vedotin, or DX3 vedotin were examined by immunofluorescence to visualize DX3, tubulin, and DAPI nuclear counterstain. No apparent changes in cell morphology or tubulin distribution were associated with DX3 or trastuzumab vedotin compared to control. In contrast, cells treated with DX3 vedotin exhibited rounding, aggregation, membrane blebbing, and contraction with reduced cytoplasmic extensions (**Fig. 6A, B**). Notably, a mitotic cell in apparent anaphase was seen in cells treated with DX3 vedotin in which the DX3 immunostain facilitated visualization of the separating chromosomes (**Fig. 6B**). Taken together, these results indicate that DX3 vedotin retains the cell-penetrating activity of DX3 and the microtubule inhibiting function of MMAE.

**Figure 6.**
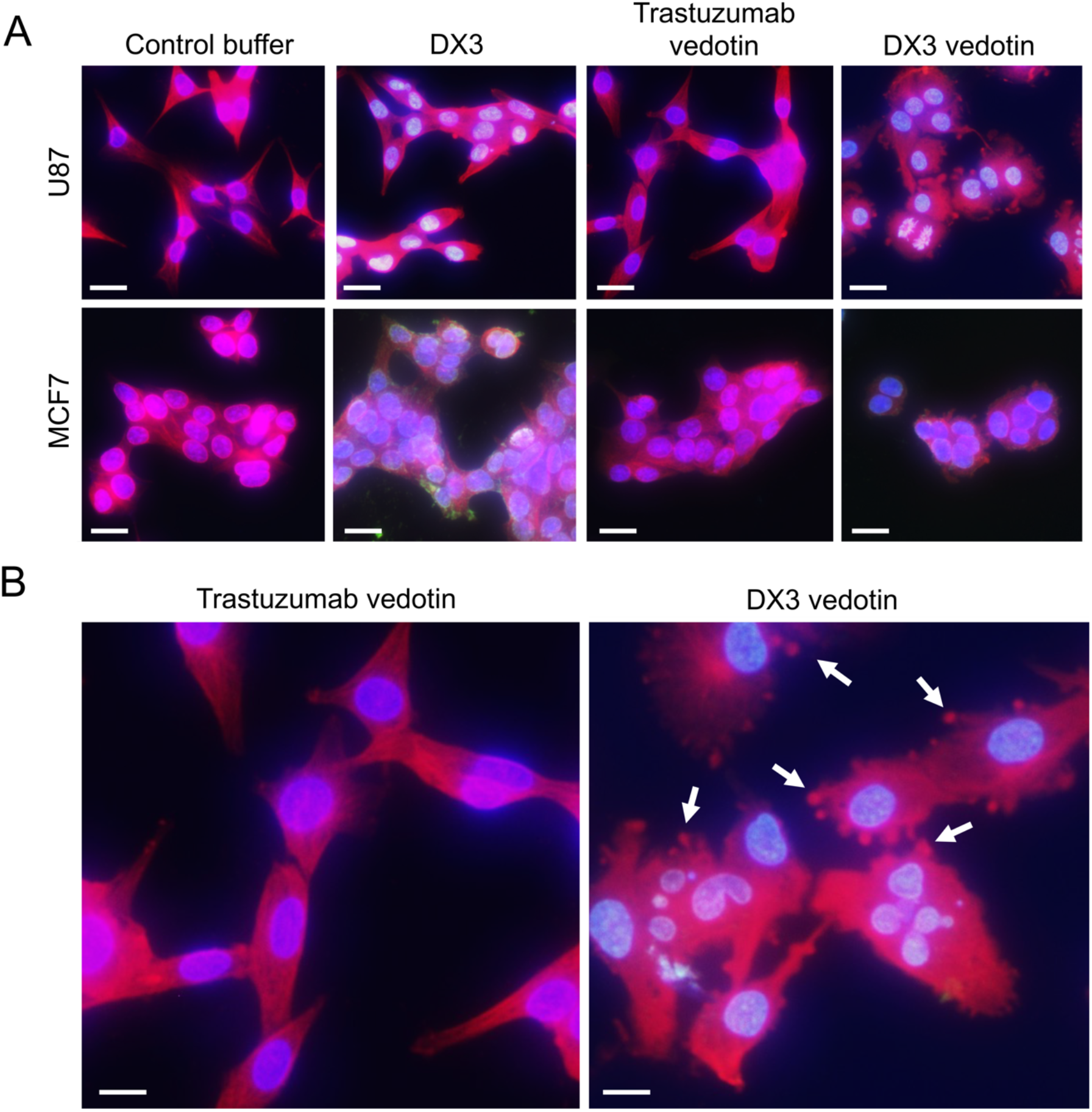
DX3 vedotin causes morphologic changes in cancer cells. (**A and B**) Merged images of DAPI, AF488 anti-IgG, and AF555 anti-tubulin stained U87 and MCF7 cells treated with control or 3 μM free DX3, trastuzumab vedotin, or DX3 vedotin are shown in (**A**) and higher magnification views of trastuzumab vedotin and DX3 vedotin-treated U87 cells in (**B**). DX3 vedotin caused a loss of cell contours and induced membrane blebbing, most evident in the U87 cells. In contrast, free DX3 and trastuzumab vedotin did not cause any apparent morphologic changes compared to control. Arrows in (**B**) denote examples of membrane blebbing. Bars: 30 μm (**A**) and 15 μm (**B**).

### DX3 vedotin is toxic to MCF7 tumors in vivo

DX3 vedotin efficacy against an HR-proficient malignancy was evaluated in an MCF7 breast cancer model. Cultured MCF7 cells were treated with serial dilutions of isotype control vedotin, DX3 vedotin, or free DX3 for five days, with cytotoxicity subsequently quantified by CellTiterGlo. Consistent with the morphology studies shown in **Fig. 6**, isotype control and DX3 vedotin yielded IC50 of 73.14 nM and 11.6 nM, respectively, while free DX3 had minimal effect on the cells (**Fig. 7A**).

**Figure 7.**
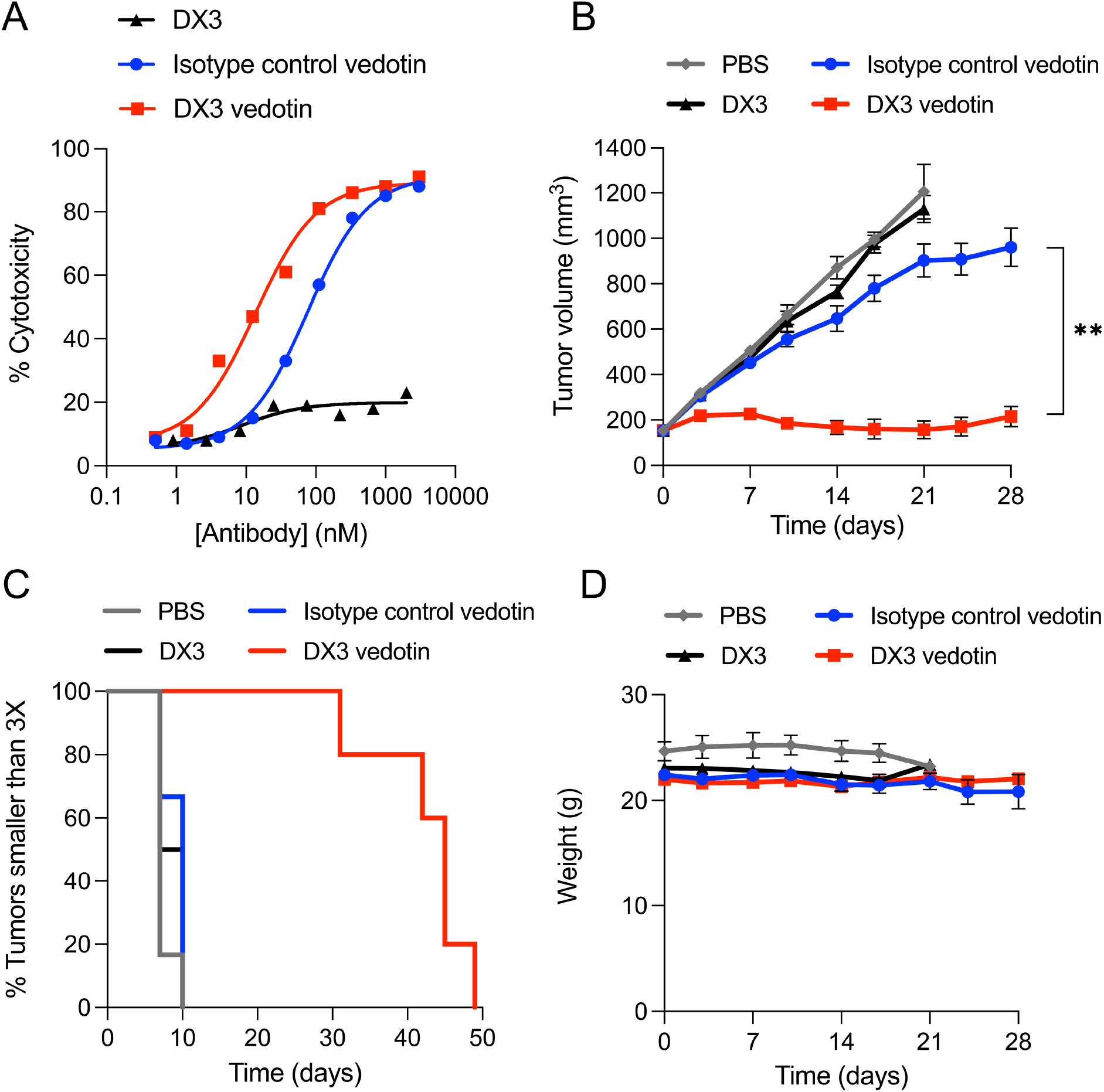
DX3 vedotin is toxic to MCF7 cancer cells and tumors. (**A**) DX3 vedotin is toxic to MCF7 cancer cells. Percent cytotoxicity in MCF7 cells after a five-day incubation with serial dilutions of DX3, DX3 vedotin, or isotype control vedotin was determined using CellTiterGlo. Respective IC50s for DX3 vedotin, isotype control vedotin, and DX3 alone were 11.6 nM, 73.14 nM, and not reached. (**B-D**) DX3 vedotin suppresses MCF7 tumor growth *in vivo*. Mice bearing MCF7 subcutaneous xenograft flank tumors with mean volume ∼150 mm^3^ were randomized to treatment with IP injection of control PBS, isotype control vedotin (10 mg/kg), DX3 vedotin (10 mg/kg), or unconjugated DX3 (10 mg/kg) once a week for four weeks (treatments were on days 0, 7, 14, and 21 in the figures). DX3 vedotin showed the greatest inhibition of tumors compared to all controls, with mean tumor volumes one week after completion of treatment 216±45 and 960±85 mm^3^ in mice treated with DX3 vedotin or isotype control vedotin, respectively (P=0.004, two-tailed Student’s t-test) (**B**). Median tumor tripling times associated with PBS, DX3, or isotype control vedotin were 7, 8.5, and 10 days, respectively, compared to 45 days with DX3 vedotin (P≤0.002 compared to all other groups, log-rank test). Kaplan-Meier plots are shown in (**C**). (**D**) All treatments were well tolerated based on behavior and body mass.

Mice bearing MCF7 subcutaneous xenograft flank tumors with mean volume ∼150 mm^3^ were randomized to groups for treatment with intraperitoneal (IP) injection of control PBS, isotype control vedotin (10 mg/kg), DX3 vedotin (10 mg/kg), or free DX3 (10 mg/kg) (N=6 mice per group) once a week for four weeks. Study endpoints were tumor volume or significant changes in behavior or weights. DX3 vedotin caused significant suppression of tumor growth, with mean tumor volumes one week after completion of treatment of 216±45 and 960±85 mm^3^ in mice treated with DX3 vedotin or isotype control vedotin, respectively (P=0.004, two-tailed Student’s t-test) (**Fig. 7B**). Median tumor tripling time in mice treated with PBS, DX3, or isotype control vedotin was 7, 8.5, and 10 days, respectively, compared to 45 days in mice treated with DX3 vedotin (P≤0.002 compared to all controls, log-rank test) (**Fig. 7C**). The experiment was stopped at day 60, selected based on lifespan of the implanted 17-β-estradiol pellets. All treatments were well-tolerated, with no significant differences in body mass observed throughout the study (**Fig. 7D**).

## Discussion

This work establishes proof-of-concept for DNA-targeting Deoxymab-based ADCs. The approach is based on DNA release by necrotic tumors and concurrent nucleoside salvage by live cancer cells promoting Deoxymab localization and uptake into tumors. In contrast to cell surface antigens that are depleted during therapy, tumor cell turnover and death yields a continuously renewing source of the DNA antigen that draws Deoxymab to tumor microenvironments. This strategy was previously demonstrated in work with 3E10-decorated drug-loaded nanocarriers wherein treatment that increased extracellular DNA in tumor environments enhanced tumor targeting by 3E10 formulations (8). However, the size of nanocarriers is expected to interfere with membrane transit by the Deoxymabs. In the present study, DX3 vedotin was well tolerated and highly effective in suppressing MCF7 breast cancer tumors. This finding demonstrates that DNA-targeting and cell-penetrating ADCs based have the potential to be used against tumors lacking specifically targetable surface antigens. Further, whereas single agent Deoxymabs have activity against DNA repair-deficient tumors, this work expands the potential for Deoxymabs to be used as a component of ADCs to treat malignancies with intact DNA repair mechanisms.

DNA is not unique to malignant tissues, which raises the possibility for off-target localization of Deoxymab ADCs. However, the Deoxymab ADC technology relies on the relative abundance of DNA in tumor microenvironments compared to normal tissues. In the present study, there was no apparent increase in toxicity of DX3 vedotin compared to any controls in the MCF7 tumor study. We believe this is due to preferential accumulation of DX3 vedotin in tumors rather than normal tissues. In addition, the greater efficacy of DX3 vedotin compared to isotype control vedotin *in vivo* suggests that DNA-targeting by DX3 enhances ADC tumor bioavailability beyond what is already conferred by the enhanced permeability and retention effect associated with solid tumors (14).

Deoxymabs also have potential to be used for targeting ADCs to regions other than tumor, such as sites of tissue injury where DNA is released by damaged cells related to underlying genetic conditions, ischemic insults, or trauma. The DNA-targeting activity of 3E10 has been leveraged to deliver linked chaperone proteins to ischemic brain and regions of myocardial infarctions, and Deoxymab ADCs are likely to exhibit similar distribution to such sites (15, 16). Moreover, the recent report that DX1 crosses the blood-brain barrier in an ENT2-dependent manner (9) raises the possibility that Deoxymabs may be able to ferry cargo drugs into the central nervous system. Overall, Deoxymabs are compelling agents for use in oncology as single agents against DNA repair-deficient tumors, as delivery ligands for nanocarriers or linked proteins, and now as components of cell-penetrating ADCs for targeting regions enriched in DNA.

## Methods

### Experimental antibodies

DX3 (PAT-DX3, Patrys Ltd., Melbourne, Australia) was generated in CHO cells and purified as previously described (9). Trastuzumab and isotype control IgG used for generation of control ADCs were provided by Syngene International Ltd. and WuXi Biologics, respectively. IgG control for the cell-penetration assay was obtained from Leinco technologies (I-118).

### Cell lines

MCF7 and U87 cells were obtained from ATCC (Manassas, VA), which confirms identity and mycoplasma-free status by STR profile and RT-PCR. BRCA2-proficient and deficient DLD1 cells were obtained from Horizon Discovery (Cambridge, UK). Cells were used within 6 months of receipt or resuscitation.

### DNA binding assay

KDs for DX3-DNA binding were determined by surface plasmon resonance using the Biacore T200 (Cytiva, Marlborough, MA, USA). The CM5 sensor chip in flow cells (FC) 1 and 2 were coated with neutravidin using the Amine Coupling Kit (Cytiva, BR100050). FC1 was used as reference cell. A 30-mer biotinylated single-stranded DNA ligand (0.025 μM, Sigma-Aldrich) was then immobilized onto the MC5 chip in FC2 through interaction with neutravidin. FC2 was used as the active flow cell. After three conditioning cycles with 5 M NaCl and six startup cycles with PBS pH 7.4, the analyte DX3 in PBS was added with flow rote 30 μl/min, contact time 120 seconds, and dissociation time 400 seconds. Results were analyzed using BIA Evaluation Software version 3.1 using a 1:1 fit kinetic binding analysis. The experiment was performed twice, with KDs of 113 nM and 112 nM determined in each experiment.

### DX3 cell penetration assays

DLD1, MCF7, or U87 cells were treated with the specified doses of control IgG (Leinco Technologies, I-118) or DX3 for one hour, after which cells were washed, fixed in chilled 100% ethanol, and immunostained to detect antibody with Alexa Fluor 488 (AF488) conjugated secondary antibody detection and DAPI counterstain as previously described (5, 6, 9). Images were obtained using an EVOS fl digital fluorescence microscope (Advanced Microscopy Group, Bothell, WA).

### DLD1 colon cancer xenograft study

Female nude mice were injected subcutaneously with matched DLD1 WT and BRCA2-deficient cells to concurrently generate HR-proficient and deficient tumors in the same mouse. 2×10^6^ DLD1 WT cells were injected in the left flank and 2×10^6^ DLD1 BRCA2-deficient cells in the right flank. Once tumors reached mean volume of ∼100 mm^3^ mice were randomized to groups for tail vein administration of vehicle control (N=8) or DX3 (50 mg/kg) (N=8) twice per week. Tumor volumes were tracked by caliper measurements and mouse weights and behaviors monitored through the study. WT tumors were observed to grow faster than the BRCA2-deficient tumors and therefore dictated timing of endpoint for tumor size. Kaplan-Meier plots of tumor tripling time (relative to tumor volumes on day 0 of treatment) were generated in Prism 9.5.1 (GraphPad Software, San Diego, CA, USA). Studies were conducted under an IACUC approved protocol, and mice were humanely euthanized for endpoints of tumor size, significant weight loss, or behavior changes.

### Generation of DX3 vedotin and control ADCs trastuzumab vedotin and isotype control vedotin

Trastuzumab, isotype control IgG, or DX3 at 5 mg/mL were incubated with TCEP at 37°C for two hours for reduction, followed by conjugation to vcMMAE by incubation in 50 mM PBS pH 7.0 for two hours at 4°C. DAR, purity, and percentage of free drug was determined by HIC-HPLC, SEC-HPLC, and LC-MS. ADCs were >98% pure and had undetectable free drug. Minimal levels of endotoxin were confirmed by PTS Cartridge.

### ADC cell penetration assays

MCF7 and U87 cells were treated with control media or media containing 3 μM DX3, trastuzumab vedotin, or DX3 vedotin for one hour, after which cells were washed, fixed in chilled 100% ethanol, and immunostained to detect IgG with AF488-conjugated anti-IgG antibody. Cells were co-stained for β-tubulin (480011, Thermo Fisher Scientific) with signal detected by Alexa Fluor 555 (AF555) conjugated secondary antibody. DAPI nuclear counterstain was used for nuclear identification as previously described (5, 6, 9). Images were obtained using an EVOS fl digital fluorescence microscope (Advanced Microscopy Group, Bothell, WA).

### Cytotoxicity assays

MCF7 cells cultured in 96-well plates were treated with titrated doses of isotype control vedotin, DX3 vedotin, or DX3 alone for five days. Cell survival was evaluated by CellTiterGlo and plotted in Prism version 9.5.1.

### MCF7 breast cancer xenograft study

Female Balb/c nude mice were administered 60-day release 17-β-estradiol pellets, and three days later injected subcutaneously with 5×10^6^ MCF7 cells in Matrigel to generate flank tumors. Once tumors reached mean volume of ∼150 mm^3^ mice were randomized to groups for intraperitoneal treatment with control PBS (N=6), isotype control vedotin (10 mg/kg) (N=6), DX3 vedotin (10 mg/kg) (N=6), or DX3 alone (10 mg/kg) (N=6) once a week for four weeks. Tumor volumes were tracked by caliper measurements and mouse weights and behaviors monitored through the study. Kaplan-Meier plots of tumor tripling time (relative to tumor volumes on day 0 of treatment) were generated in Prism 9.5.1 (GraphPad Software, San Diego, CA, USA). One mouse in the DX3 vedotin group was lost (unrelated to tumor or treatment) prior to reaching tumor tripling time and was excluded from the Kaplan-Meier analysis. Studies were conducted under an IACUC-approved protocol, and mice were humanely euthanized for endpoints of tumor size, significant weight loss, or behavior changes.

### Statistics

P values were determined by two-tailed Student’s t-test or log-rank test in Prism 9.5.1. P≤0.05 was considered significant.

## Acknowledgements

This work was supported in part by a Patrys Ltd. sponsored research agreement and the Yale Department of Therapeutic Radiology (JEH).

## Conflict of Interest

AE has equity interest in and receives consulting fees and equity options from Patrys Ltd. VD and JAC are employees of and have equity interest in Patrys Ltd. JEH has related intellectual property (IP) and equity interest in and receives grant support and consulting fees in the form of equity options from Patrys Ltd., a biotechnology company that has licensed IP that is the basis of this work. The remaining authors declare no conflicts of interest.

